# Sirtuin 5 levels are limiting in preserving cardiac function and suppressing fibrosis in response to pressure overload

**DOI:** 10.1101/2021.06.15.448619

**Authors:** Angela H. Guo, Rachael K. Baliira, Mary E. Skinner, Surinder Kumar, Anthony Andren, Li Zhang, Shaday Michan, Norma J. Davis, Merissa W. Maccani, Sharlene M. Day, David A. Sinclair, Costas A. Lyssiotis, Adam B. Stein, David B. Lombard

**Author notes:** Correspondence: University of Michigan, 3015 BSRB, 109 Zina Pitcher Place, Ann Arbor, MI 48109.

## Abstract

Heart failure (HF) is defined as an inability of the heart to pump blood adequately to meet the body’s metabolic demands. HF with reduced systolic function is characterized by cardiac hypertrophy, ventricular fibrosis and remodeling, and decreased cardiac contractility, leading to cardiac functional impairment and death. Transverse aortic constriction (TAC) is a well-established model for inducing hypertrophy and HF in rodents. Mice globally deficient in sirtuin 5 (SIRT5), a NAD^+^-dependent deacylase, are hypersensitive to cardiac stress and display increased mortality after TAC. Prior studies assessing SIRT5 functions in the heart have all employed loss-of-function approaches. In this study, we generated SIRT5 overexpressing (SIRT5OE) mice, and evaluated their response to chronic pressure overload induced by TAC. Compared to littermate controls, SIRT5OE mice were protected from left ventricular dilation and impaired ejection fraction, adverse functional consequences of TAC. Transcriptomic analyses revealed that SIRT5 suppresses key HF sequelae, including the metabolic switch from fatty acid oxidation to glycolysis, immune activation, and increased fibrotic signaling. We conclude that SIRT5 is a limiting factor in the preservation of cardiac function in response to experimental pressure overload.

## Introduction

Heart failure (HF) is the convergent final pathway of a variety of cardiovascular pathologies that leads to insufficient oxygen delivery to meet the metabolic demands of the body. The incidence of HF is predicted to increase by 46% between 2012 and 2030 in the US, mainly due to an increase in human lifespan coupled with improved survival rates following cardiovascular injury (1). In 2018 alone, HF claimed 83,616 lives (2). HF with systolic dysfunction is driven by a progressive decline in contractile function and chronic hemodynamic overload, and characterized by ventricular hypertrophy and remodeling, neurohormonal compensation mechanisms, and myocardial damage. Key cellular mechanisms thought to contribute to HF include myocyte death, reduced ability to maintain calcium homeostasis, and changes in production and utilization of high-energy phosphates (3). However, the mechanistic changes that lead to HF are still incompletely understood, and currently there are no mechanism-based treatments for HF.

Transverse aortic constriction (TAC) is a well-established experimental rodent model for pressure overload-induced HF. In TAC, the aorta is surgically narrowed, chronically increasing resistance to outflow from the left ventricle (LV) (4). This increased left ventricular burden initially induces concentric hypertrophy to compensate for TAC-induced elevated resistance. However, chronic pressure overload leads to contractile dysfunction and extensive remodeling, including fibrosis and ventricular dilation, resulting in functional decline. Over time, the deteriorating LV becomes unable to meet the hemodynamic needs of the body, resulting in progressive HF (5).

Sirtuins are a family of NAD^+^-dependent deacylases, ADP-ribosyltransferases, and lipoamidases involved in cellular stress responses. In mammals, each of the seven sirtuins possesses a distinct combination of enzymatic activities, molecular targets, subcellular localization, and tissue expression (6). Sirtuin 5 (SIRT5) possesses robust deacylase activity against lysine succinylation, malonylation, and glutarylation, negatively-charged lysine acyl modifications derived from their respective Coenzyme A (CoA) species (7–11). SIRT5 target proteins reside in the mitochondrial matrix, cytosol, and nucleus; many function in metabolic pathways, including the citric acid cycle (*e.g.* pyruvate dehydrogenase and succinate dehydrogenase (SDH)) and fatty acid oxidation (*e.g.* trifunctional enzyme subunit alpha (ECHA)) (8, 12). Although many SIRT5 targets take part in key metabolic pathways, no major metabolic defects have been observed in *Sirt5* knockout (SIRT5KO) or overexpressing (OE) mice under basal, unstressed conditions (13–15).

Among normal tissues, a major site of SIRT5 function appears to be the heart. SIRT5KO mice display a modest, age-associated decrease in cardiac function, along with increased fibrosis and hypertrophy (12). SIRT5KO mice show marked hypersensitivity to cardiac stress. SIRT5KO mice are more susceptible to ischemia-reperfusion (I/R) injury, showing increased infarct size and impaired recovery (16). In these studies, proteomics identified ECHA and SDH, respectively, as substrates most relevant to SIRT5’s cardioprotective activity. In response to TAC, wholebody SIRT5KOs developed more severe cardiac dysfunction, and exacerbated hypertrophy and fibrosis compared to wild-type (WT) littermates. The authors of this study propose that, during TAC, SIRT5 deficiency is associated with impaired electron flow in oxidative phosphorylation, elevated mitochondrial NADH levels, and impaired oxidative metabolism, with greater reliance on glycolysis (17). In contrast, a subsequent study by the same group using a cardiomyocyte (CM)-specific SIRT5KO strain showed no differences between WT and SIRT5 mutants in response to TAC (18).

In this study, we generated a novel strain of SIRT5 overexpressing (SIRT5OE) mice and evaluated their response to cardiac pressure overload. SIRT5OE mice displayed no obvious cardiac phenotype and very few gene expression changes in the absence of stress compared to WT littermates. However, SIRT5OE mice showed robust protection against TAC-induced LV dilation and subsequent functional decline. Transcriptomic analysis linked the protection conferred by SIRT5 overexpression to the suppression of key transcriptional and signaling events accompanying ventricular hypertrophy and HF, including cardiac fibrosis, inflammatory cytokine signaling, and the metabolic switch from fatty acid oxidation to glycolysis. Our results suggest that SIRT5 levels are limiting in the context of the cardiac response to pressure overload.

## Results

### Generation and characterization of SIRT5-overexpressing mice

To investigate the effects of increased SIRT5 on cardiac function, we generated a transgenic SIRT5 overexpressing (SIRT5OE) mouse strain by inserting a LoxP-STOP-LoxP-SIRT5 cassette into the 3’UTR of the *Col1A1* locus, using a well-characterized system permitting transgene insertion at a specific locus (19). SIRT5 expression is driven by the CAGGS promoter (Supplemental Figure 1A) (20). SIRT5OE mice were born at normal Mendelian and sex ratios, and grossly indistinguishable from their WT littermates, with no obvious differences in weight gain with age in either sex (Supplemental Figure 1B, Figure 1A). We tested the molecular effects of SIRT5 overexpression on bulk protein acylation in the heart and found modest, nonsignificant decreases in total lysine succinylation (Ksucc) and lysine malonylation (Kmal) (Fig. 1B-D).

**Figure 1.**
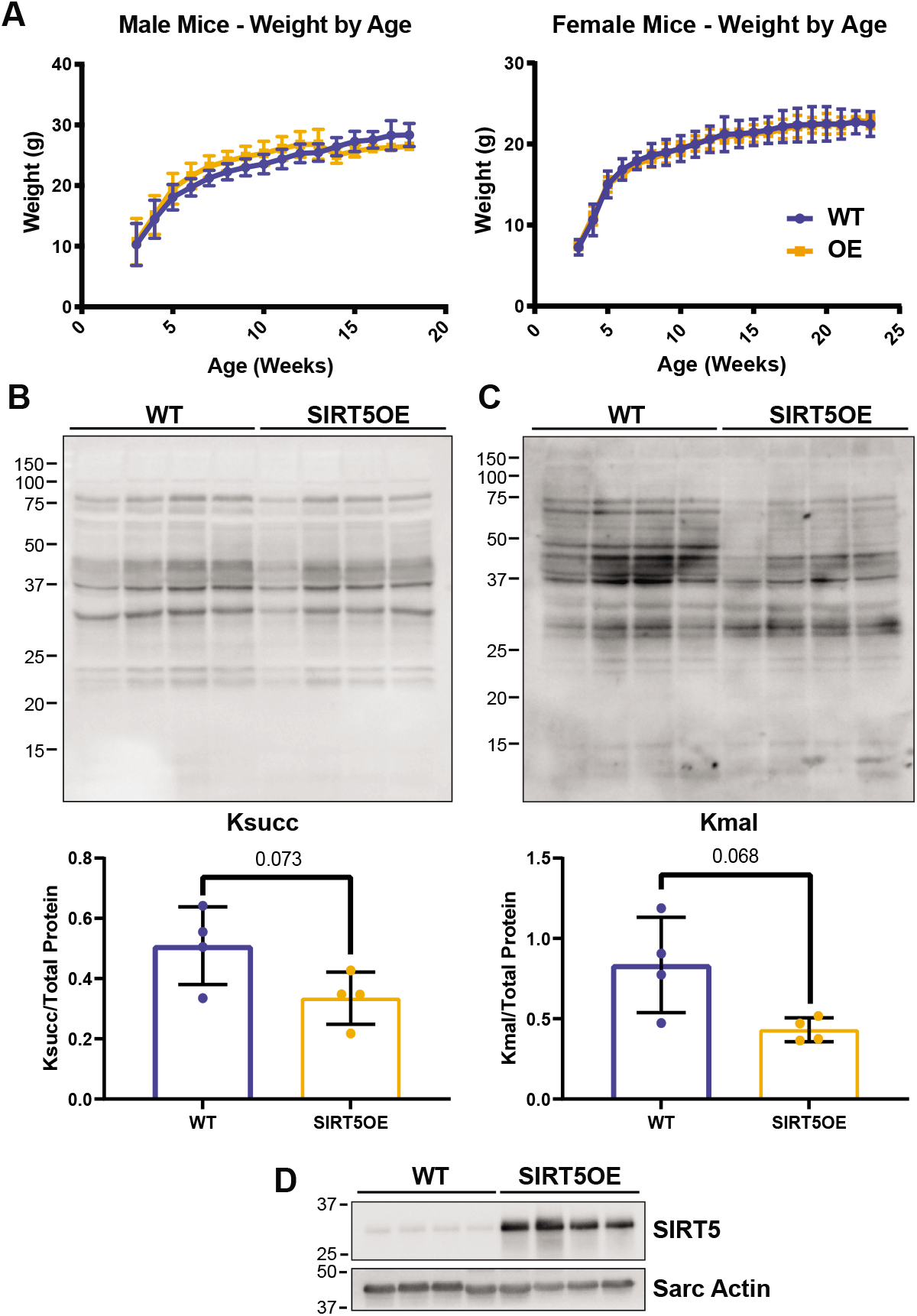
SIRT5OE mice exhibit slightly decreased cardiac lysine acylation but no gross phenotypic changes. A, weight (g) of WT and SIRT5OE male and female mice, measured with age. B-C, representative western blot analysis of succinyl-lysine (Ksucc), and malonyl-lysine (Kmal) levels in WT and SIRTOE hearts, with quantification. D, immunoblot analysis of SIRT5 expression in samples in B-C.

### SIRT5 OE does not alter cardiac hypertrophy after pressure overload

To evaluate their sensitivity to cardiac stress, male WT and SIRT5OE littermates between 4-8 months of age were randomly assigned to sham or TAC operation (Figure 2A) (21). The TAC procedure surgically narrows the transverse aorta, generating a pressure gradient that reflects the severity of pressure overload on the heart (Figure 2B). Sham animals undergo an identical surgical procedure but without aortic banding. We first assessed whether TAC changed total Ksucc levels in the heart. Immunoblot showed that SIRT5OE caused a modest decrease in total Ksucc levels. Additionally, sustained pressure overload led to reductions in total Ksucc levels regardless of genotype, mirroring the results observed in SIRT5KO sham and TAC mice (Figure 2C-D) (17). To complement these data, we obtained heart tissue samples from human HF patients. Similar to the mouse hearts, Ksucc levels trended lower in the human failing hearts compared to the control samples (p=0.07) (Figure 2E-F). Therefore, cardiac stress leads to decreased levels of Ksucc, independent of *SIRT5* genotype, in both mouse and human samples.

**Figure 2.**
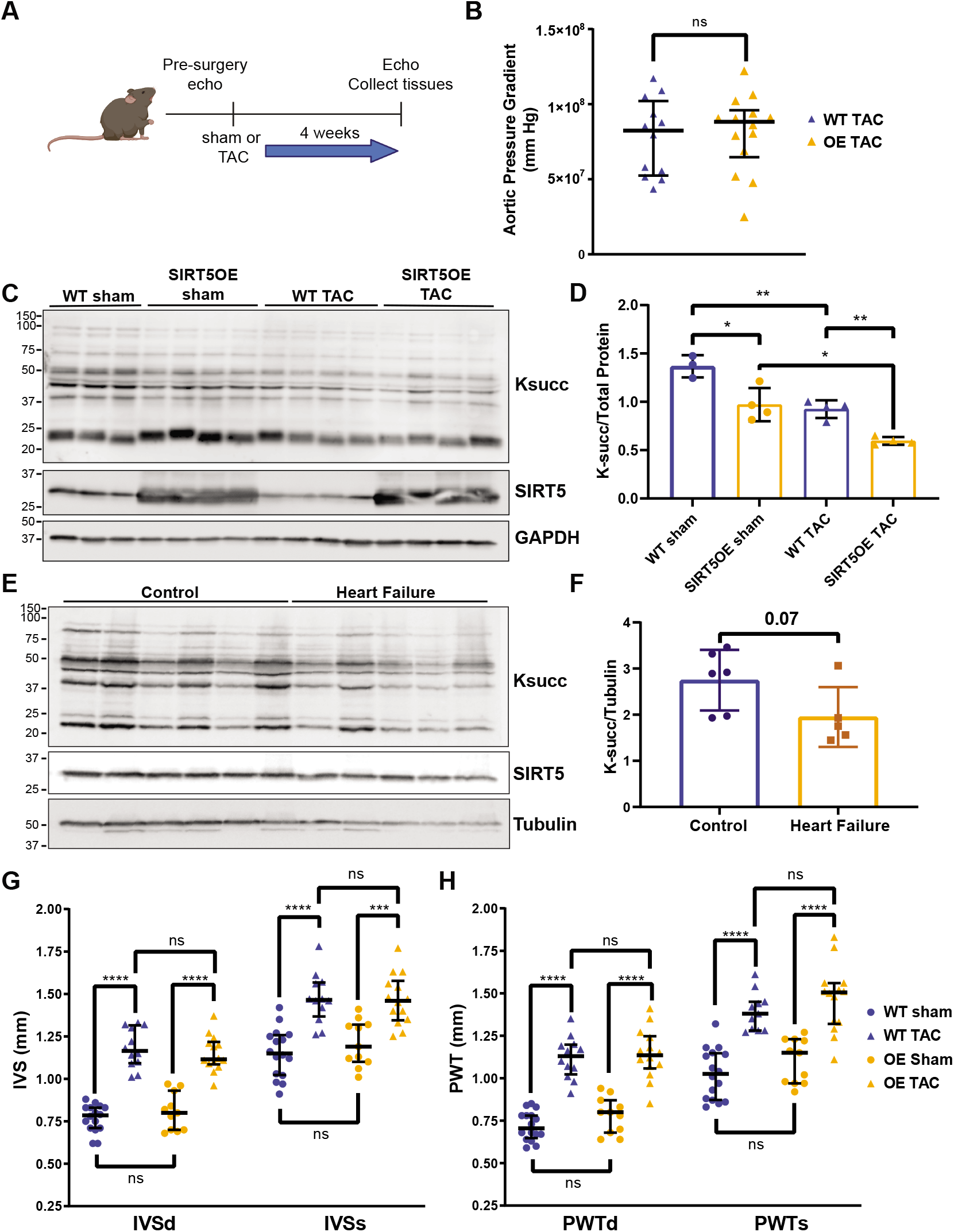
WT and SIRT5OE mice develop similar cardiac hypertrophy 4 weeks after TAC. Echocardiography was performed on WT sham (n=16), WT TAC (n=12), SIRT5OE sham (n=10) and SIRT5OE TAC (n=14) mice to assess changes in physiology prior to and four weeks after surgery. A, depiction of groups, procedures, and timeline of surgery. B, aortic pressure gradient in mice after TAC. C-D, immunoblot analysis of Ksucc in sham and TAC mice with quantification. E-F, immunoblot analysis of Ksucc with quantification in heart tissue from patients with quantification. G, diastolic and systolic measurement of interventricular septum (IVS) thickness. H, diastolic and systolic measurement of posterior wall thickness (PWT). Statistical significance was determined using Student’s T-test for 2-group analysis or two-way ANOVA followed by Sidak’s correction for multiple comparisons for 4-group analyses.

To assess cardiac function, echocardiograms (echos) were performed prior to surgery, and then repeated at the end of a 4-week observational period to assess cardiac structure and function (Table 1, Figure 2A). WT and SIRT5OE sham animals did not exhibit any notable differences by echo, indicating that global SIRT5 overexpression does not alter baseline cardiac structure or function (Table 1). Additionally, the magnitude of the aortic pressure gradient was comparable between WT and SIRT5OE TAC mice (Figure 2B). Thus, any differences observed between genotypes following TAC were not due to discrepancies in the degree of cardiac outflow obstruction. Concentric cardiac hypertrophy is a temporary compensatory mechanism developed to relieve the stress induced by pressure overload (22). Echo measurement of the interventricular septum (IVS) and the posterior wall (PW) thickness during both systole and diastole showed comparable LV hypertrophy 4 weeks after TAC in all mice irrespective of genotype (Figure 2G, H). Therefore, SIRT5 overexpression does not affect cardiac concentric hypertrophy in response to TAC.

**Table 1.** Echocardiogram measurements 4 weeks after surgery for WT, SIRT5 sham and TAC mice. Values listed as mean ± SEM. Statistical significance was determined using two-way ANOVA. Significance markers: (*) p < 0.05, (+) p < 0.1.

| Table 1. Echocardiogram measurements 4 weeks after surgery for WT, SIRT5 sham and TAC mice |  |  |  |  |  |  |  |
| --- | --- | --- | --- | --- | --- | --- | --- |
|  | WT sham<br>(n=16) | WT TAC<br>(n=12) | SIRT5OE<br>sham<br>(n=11) | SIRT5OE TAC<br>(n=14) | Genotype<br>Effect | TAC<br>Effect | Interaction<br>Effect |
| Age at Procedure (months) | 5.32±0.60 | 5.26±0.50 | 5.88±0.82 | 5.26±0.46 |  |  |  |
| Heart rate (bpm) | 504.75±13.31 | 487.17±19.47 | 502.26±17.59 | 452.04±14.58 | 0.250 | 0.041* | 0.317 |
| LV Mass/BW | 3.86±0.11 | 9.37±0.56 | 4.01±0.13 | 7.55±0.32 | 0.012* | <0.001* | 0.003* |
| IVS thickness diastole (mm) | 0.77±0.02 | 1.18±0.03 | 0.80±0.03 | 1.15±0.03 | 0.937 | <0.001* | 0.217 |
| IVS thickness systole (mm) | 1.15±0.04 | 1.47±0.04 | 1.21±0.04 | 1.47±0.04 | 0.540 | <0.001* | 0.433 |
| LV diameter diastole (mm) | 3.89±0.05 | 4.61±0.14 | 3.82±0.10 | 4.11±0.08 | 0.004* | <0.001* | 0.027* |
| LV diameter systole (mm) | 2.86±0.05 | 3.81±0.15 | 2.72±0.10 | 3.14±0.09 | <0.001* | <0.001* | 0.007* |
| PW thickness diastole (mm) | 0.72±0.02 | 1.12±0.04 | 0.78±0.03 | 1.14±0.04 | 0.208 | <0.001* | 0.523 |
| PW thickness systole (mm) | 1.02±0.04 | 1.35±0.05 | 1.10±0.04 | 1.47±0.05 | 0.082+ | <0.001* | 0.993 |
| LV Volume diastole (uL) | 65.76±2.11 | 99.41±7.53 | 63.33±3.74 | 75.44±3.51 | 0.004* | <0.001* | 0.018* |
| LV Volume systole (uL) | 31.35±1.39 | 63.99±6.57 | 29.05±2.65 | 39.86±2.70 | <0.001* | <0.001* | 0.004* |
| Ejection Fraction (%) | 0.52±0.01 | 0.36±0.02 | 0.55±0.02 | 0.47±0.02 | 0.002* | <0.001* | 0.036* |
| Fractional Shortening (%) | 0.27±0.01 | 0.18±0.01 | 0.28±0.02 | 0.24±0.01 | 0.003* | <0.001* | 0.061+ |

### SIRT5 overexpression protects against TAC-induced heart failure

After prolonged pressure overload, concentric hypertrophy progresses to ventricular dilation and heart failure (HF) (5). To determine if SIRT5OE alters this progression, we measured LV diameter by echo. After four weeks of TAC, the LVs of WT mice showed significantly increased diameter, indicating that these mice had transitioned from adaptive to maladaptive ventricular hypertrophy. In comparison, LV diameter did not significantly increase in SIRT5OE TAC mice (Figure 3A). The combination of ventricular hypertrophy and dilation was also reflected in the LV size normalized to body weight. Both genotypes showed increased normalized LV mass after TAC; however, this increase was blunted in SIRT5OE animals (Figure 3B). Ventricular dilation often leads to decreased ejection fraction (EF), a measure of systolic function. EF was significantly reduced in response to chronic pressure overload in the WT mice but preserved in the SIRT5OE mice (Figure 3C). Representative echo images of WT TAC and SIRT5OE TAC hearts are shown in Figure 3D.

**Figure 3.**
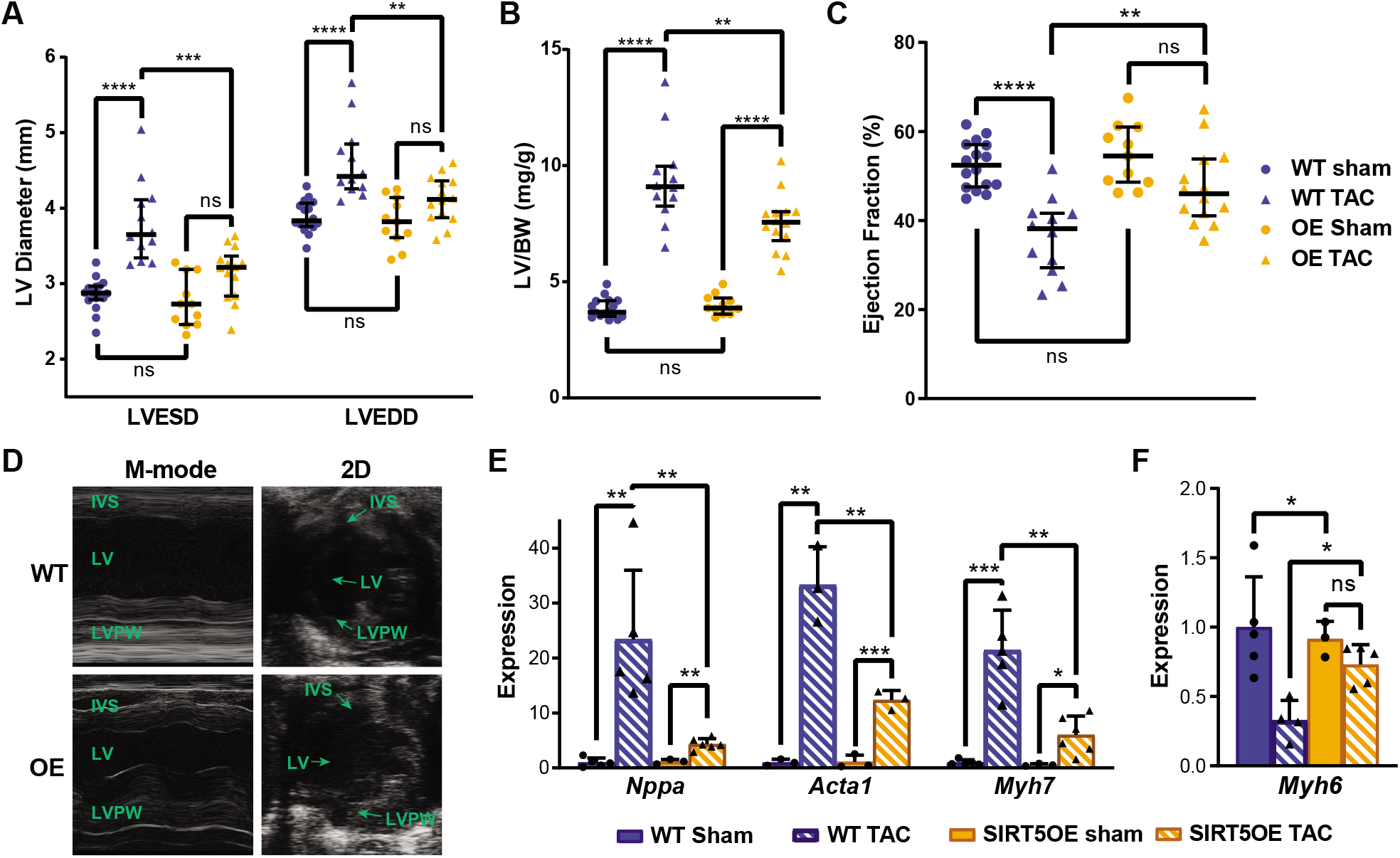
SIRT5OE mice are protected against TAC-induced heart failure. A, echo measurements of left ventricular end systolic and diastolic diameter (ESD or EDD respectively). Echocardiography was performed on WT sham (n=16), WT TAC (n=12), SIRT5OE sham (n=10) and SIRT5OE TAC (n=14). B, left ventricle mass normalized to body weight. C, ejection fraction. D, representative M-mode and 2D echo results from WT and OE mice 4 weeks after TAC. E, qRT-PCR for Nppa, Acta1, Myh6, and Myh7 expression normalized to GAPDH. Statistical significa3n8ce was determined using two-way ANOVA followed by Sidak’s correction for multiple comparisons.

To complement echo-based measurements, we assessed RNA levels of standard markers of cardiac hypertrophy and HF, atrial natriuretic peptide (*Nppa*) and skeletal muscle Actin (*Acta1*). As expected, expression of both genes increased in response to TAC, but was blunted in SIRT5OE TAC mice compared to WT TAC (Figure 3E). The failing myocardium also shifts contractile protein expression from myosin heavy chain isoform α (*Myh6*) to β (*Myh7*) (Figure 3E-F) (23). The decrease in *Myh6* expression by TAC was attenuated in SIRT5OE mice compared to WT controls. Overall, we conclude from these physiological and molecular data that SIRT5OE protects against the transition from adaptive, concentric hypertrophy to maladaptive ventricular dilation and systolic dysfunction.

### SIRT5OE blunts transcriptomic changes induced by TAC

To elucidate possible mechanisms by which SIRT5OE protects against TAC-induced stress, we performed RNA-seq transcriptomic profiling on whole hearts from the four groups (WT sham, WT TAC, SIRT5OE sham, SIRT5OE TAC). Principal component analysis (PCA) showed that SIRT5 overexpression did not appreciably alter baseline gene expression in the heart, as sham mice clustered together regardless of genotype. In contrast, TAC induced a marked transcriptional response in both genotypes. Apart from one outlier, SIRT5OE TAC mice clustered between sham animals and WT TAC animals, suggesting that SIRT5OE partially blunts the overall transcriptional response to TAC (Figure 4A). Hierarchical cluster analysis (HCA) of the top 30 genes ranked by variance generated a dendrogram with two major clades (Figure 4B). With the exception of one animal, all SIRT5OE mice clustered together with WT sham mice in one clade. The second clade contained all the WT TAC mice and one SIRT5OE TAC mouse. Genes represented in the HCA plot are markers of hypertrophy/HF (*Nppa, Myh7, Serpina3n*), fibrosis signaling (*Postn, Timp1, collagen components*), and immune response (*Lcn2, Spp1, Crlf1*) (24–27). Thus, based on both PCA and HCA, SIRT5OE TAC mice are generally more transcriptionally similar to WT sham mice than WT TAC mice.

**Figure 4.**
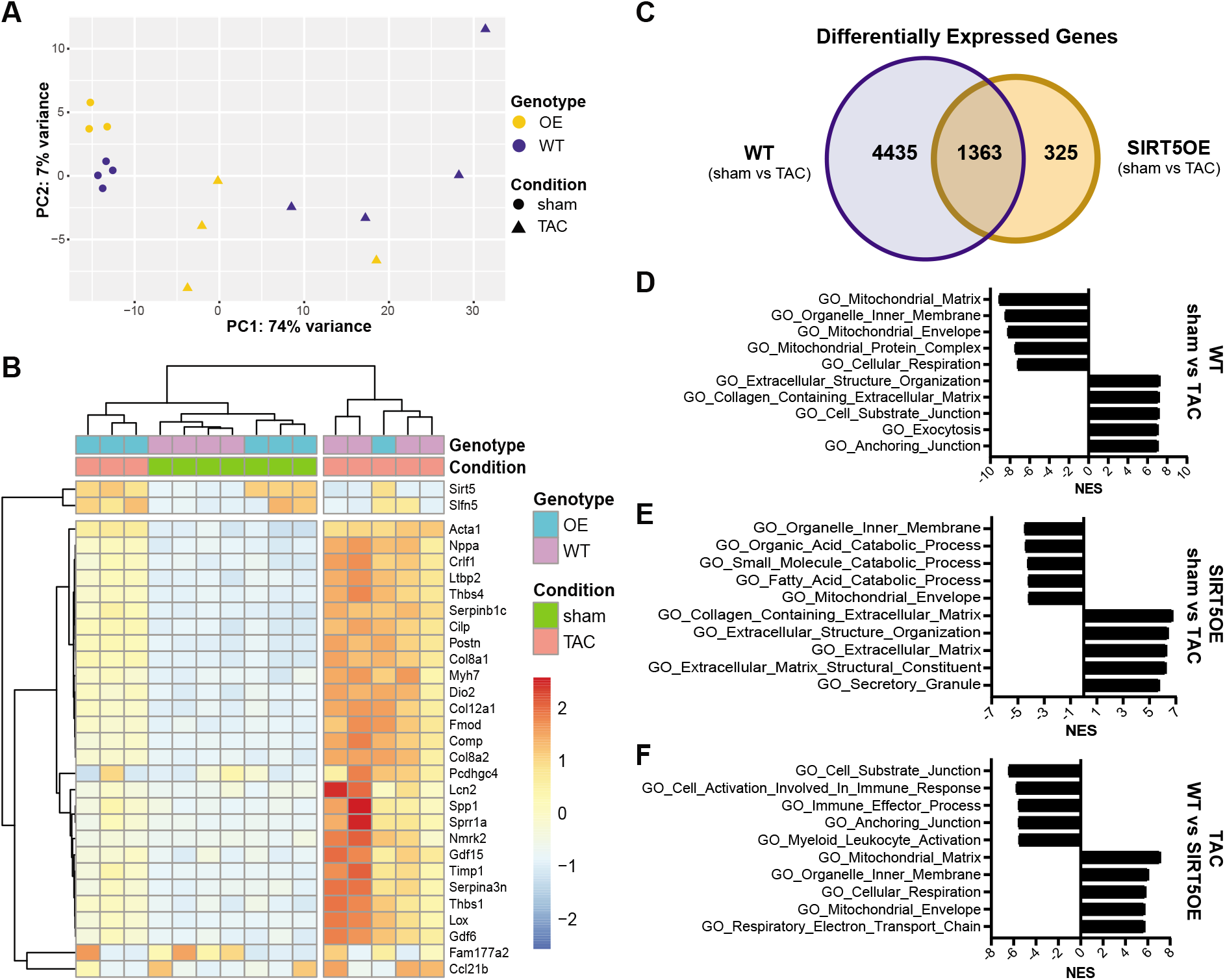
Transcriptomic analysis of heart tissue. A, principal component analysis (PCA) of RNA-sequencing data of the four groups (WT sham (n=4), WT TAC (n=4), SIRT5OE sham (n=3) and SIRT5OE TAC (n=4)). B, Hierarchical clustering analysis (HCA) of the RNA-seq data based on the top 30 genes, determined by greatest variance from the mean. Genes with higher expression compared to the mean skew red; genes with lower expression compared to the mean skew blue. C, Venn Diagram comparing the exclusive and overlapping differentially expressed genes (FDR < 0.05) in the WT (sham vs TAC) mice and the SIRT5OE (sham vs TAC) mice. D-F, Top five positively and negatively enriched Gene Ontology (GO) pathways from the indicated comparison, determined by gene set enrichment analysis (GSEA). NES: normalized enrichment score. All listed pathways were significantly enriched with a false discovery rate (FDR) < 0.0001.

We defined the criteria for differentially expressed genes (DEGs) between two groups at a false discovery rate (FDR) < 0.05. Strikingly, there were only 9 DEGs, including SIRT5, between sham WT and SIRT5OE mice, consistent with notion that SIRT5 overexpression does not significantly alter baseline cardiac biology. Following TAC, the number of DEGs increased dramatically in both genotypes. There were 4435 unique DEGs between WT sham and TAC hearts, but only 325 in the SIRT5OE sham and SIRT5OE TAC. There were 1363 DEGs shared between the WT and SIRT5OE sham to TAC comparisons, representing a core group of genes that respond to chronic pressure overload (Figure 4C). The smaller number of DEGs in SIRT5OE mice after TAC suggests that transcriptional responses induced by pressure overload are ameliorated by SIRT5 overexpression.

Unbiased analysis of gene signatures enriched by the DEGs of each group was performed using Ingenuity Pathway Analysis (IPA) and Gene Set Enrichment Analysis (GSEA) (28, 29). The top 5 most positively and negatively enriched Gene Ontology (GO) pathways based on normalized enrichment score (NES) were extracellular matrix (ECM) remodeling and mitochondrial/metabolic pathways, respectively in sham to TAC comparisons of both genotypes, albeit to different magnitudes (Figure 4D-E). The most enriched GO terms between TAC samples revealed that SIRT5OE TAC hearts show reduced activation of the immune system and suppressed ECM organization while maintaining expression of respiratory genes compared to WT TAC hearts. IPA analysis of these samples showed similar patterns of pathway enrichment (Supplemental Figure 2A-C). Table 2 lists pathways known to play critical roles in the development and progression of pathological hypertrophy and HF. All pathways are more significantly enriched by IPA in the WT sham to TAC comparison, with oxidative phosphorylation showing no enrichment in the SIRT5OE sham vs TAC analysis. In summary, based on PCA, HCA, and function enrichment analysis, the RNA-seq data support the conclusion that SIRT5 overexpression protects against pressure overload-induced HF, partially through the transcriptional suppression of molecular signaling pathways responsible for HF progression.

**Table 2.** IPA pathway analysis of cardiovascular disease related pathways in WT (sham vs TAC) and SIRT5OE (sham vs TAC) samples. All pathways are significant except for oxidative phosphorylation in SIRT5OE (sham vs TAC).

| **Table 2. IPA pathway analysis of cardiovascular disease related pathways in WT (sham vs TAC) and SIRT5OE (sham vs TAC) samples** |  |  |
| --- | --- | --- |
|  | **WT Mice**<br>**(sham vs TAC)** | **SIRT5OE Mice**<br>**(sham vs TAC)** |
| **Cardiovascular Disease-Related Pathways** | **-log(p-value)** |  |
| Oxidative Phosphorylation | 22.00 | 0.00* |
| Hepatic Fibrosis Signaling Pathway | 21.30 | 10.90 |
| Cardiac Hypertrophy Signaling (Enhanced) | 15.60 | 7.01 |
| HIF1α Signaling | 13.30 | 3.81 |
| NRF2-mediated Oxidative Stress Response | 11.70 | 3.33 |
| IL-6 Signaling | 9.50 | 3.19 |
| Renin-Angiotensin Signaling | 9.45 | 3.14 |
| Role of NFAT in Cardiac Hypertrophy | 8.58 | 4.21 |
| Angiopoietin Signaling | 8.44 | 2.61 |
| Thrombin Signaling | 7.47 | 2.69 |
| Fatty Acid β-oxidation I | 5.04 | 1.55 |
| Endothelin-1 Signaling | 4.94 | 1.79 |
| TGF-β Signaling | 4.69 | 2.07 |
| Atherosclerosis Signaling | 3.07 | 2.37 |

### Pressure overload induces similar metabolic changes in both WT and SIRT5OE hearts

Under normal conditions, the heart utilizes fatty acids as the main source of ATP generation. The failing heart increases glycolysis usage without a concomitant increase in glucose oxidation, implying an overall decrease in mitochondrial oxidative metabolism, resulting in an energy deficit (30). Gene ontology analysis revealed that WT TAC mice showed significant downregulation of pathways mapping to cellular respiration and the mitochondrial matrix. Similarly, SIRT5OE mice post-TAC also showed reduced expression of genes in various catabolic processes, including fatty acid and small molecule catabolism (Figure 4D-E). A more detailed investigation of GSEA results comparing the WT and SIRT5OE TAC transcriptomes revealed enrichment of hallmark glycolytic genes in the WT TAC samples. Conversely, genes for fatty acid metabolism and components of the respiratory electron transport chain (ETC) were enriched in the SIRT5OE TAC samples (Figure 5A). Overall, these transcriptional differences suggest that 4 weeks after surgery, SIRT5OE TAC mice show an intermediate metabolic state between healthy (sham) hearts and the failing, WT TAC hearts. These metabolic gene signatures are consistent with PCA and HCA results (Figure 4A-B).

**Figure 5.**
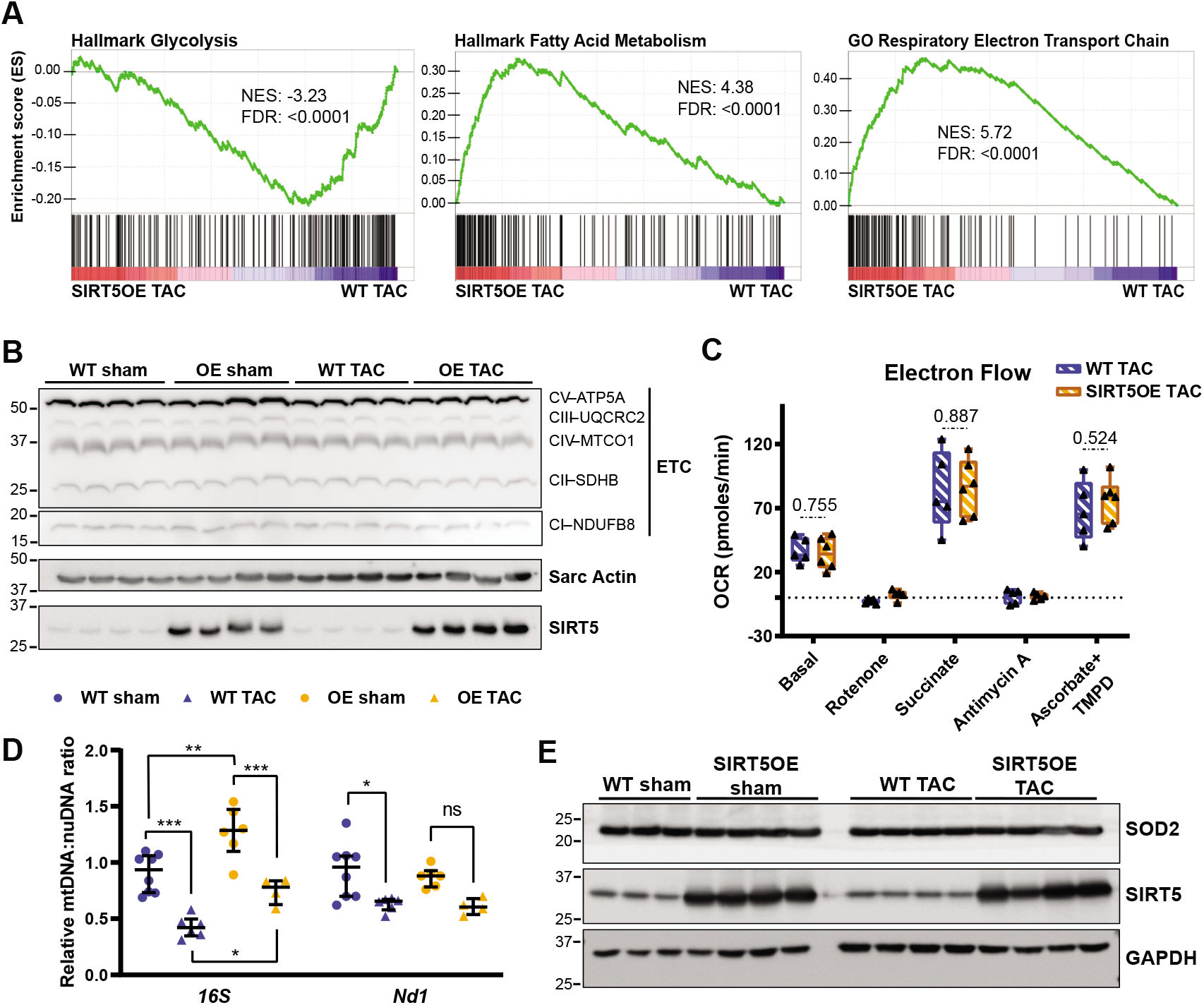
WT TAC mice exhibit a metabolic shift by RNA-seq, but are functionally comparable to SIRT5OE TAC mice. A, GSEA of SIRT5OE TAC compared to WT TAC RNA-seq samples. NES: normalized enrichment score. FDR: false discovery rate. B, immunoblot analysis of electron transport chain subunits. C, Agilent/Seahorse analysis of electron flow through the ETC in mitochondria isolated from WT (n = 5) and SIRT5OE (n = 6) hearts 4 weeks after TAC. Y-axis measures oxygen consumption rate (OCR) in pmol/minute. D, qPCR of mitochondrially encoded (mtDNA) genes 16S and ND1 normalized to HK2, encoded by the nuclear genome (nuDNA). E, SOD2 immunoblot from heart lysates.

To investigate whether RNA expression differences in these metabolic pathways also manifested at the protein level, we immunoblotted for components of the ETC but found no differences in protein expression between different genotypes or treatments, suggesting that mitochondrial respiratory complex formation are normal (Figure 5B). We then directly assessed ETC activity between WT and SIRT5OE TAC hearts using the Agilent/Seahorse XFe96 analyzer and observed similar electron flow in mitochondria isolated from hearts of each genotype after TAC (Figure 5C).

With comparable protein expression levels of the ETC and similar electron flow, we then asked whether differences in total mitochondrial content might contribute to the differences in LV function. Mitochondrial mass was quantified two ways. First, qPCR was performed on the mitochondrially-encoded *16S* and *Nd1* genes, normalized to *Hk2,* a gene encoded by the nuclear genome, and to WT sham samples (31). Second, we also performed immunoblotting for SOD2, a protein localized to the mitochondria. Neither assay revealed any major differences in total mitochondrial content between WT and SIRT5OE TAC groups (Figure 5D-E). From these data, we conclude that while WT and SIRT5OE TAC hearts show transcriptional differences related to metabolic pathways, these expression differences do not manifest at the level of electron transport chain activity, changes in levels of respiratory complexes, or mitochondrial content.

To determine if the cardio-protective properties of SIRT5 overexpression is derived from alterations in the metabolome, we performed unbiased metabolomics on cardiac samples across all four groups using LC-based mass spectrometry. PCA showed minimal variance between samples on the first two principal components (Figure 6A). TAC was the main influence on sample clustering, with minimal contribution by genotypic differences. Analysis of the data for metabolite differences between genotypes also supported this conclusion, with only 6 and 4 metabolites significantly altered for WT sham to SIRT5OE sham and WT TAC to SIRT5OE TAC comparisons, respectively. In contrast, the WT sham to TAC comparison showed the greatest number of significantly different metabolites, 44, followed by the SIRT5OE sham to TAC comparison, 25. Next, metabolite data was analyzed via MetaboAnalyst to analyze the pathways enriched across genotypes and conditions, but these analyses did not generate any insights into genotypic differences, most likely due to the limited number of significantly altered metabolites (not shown).

**Figure 6.**
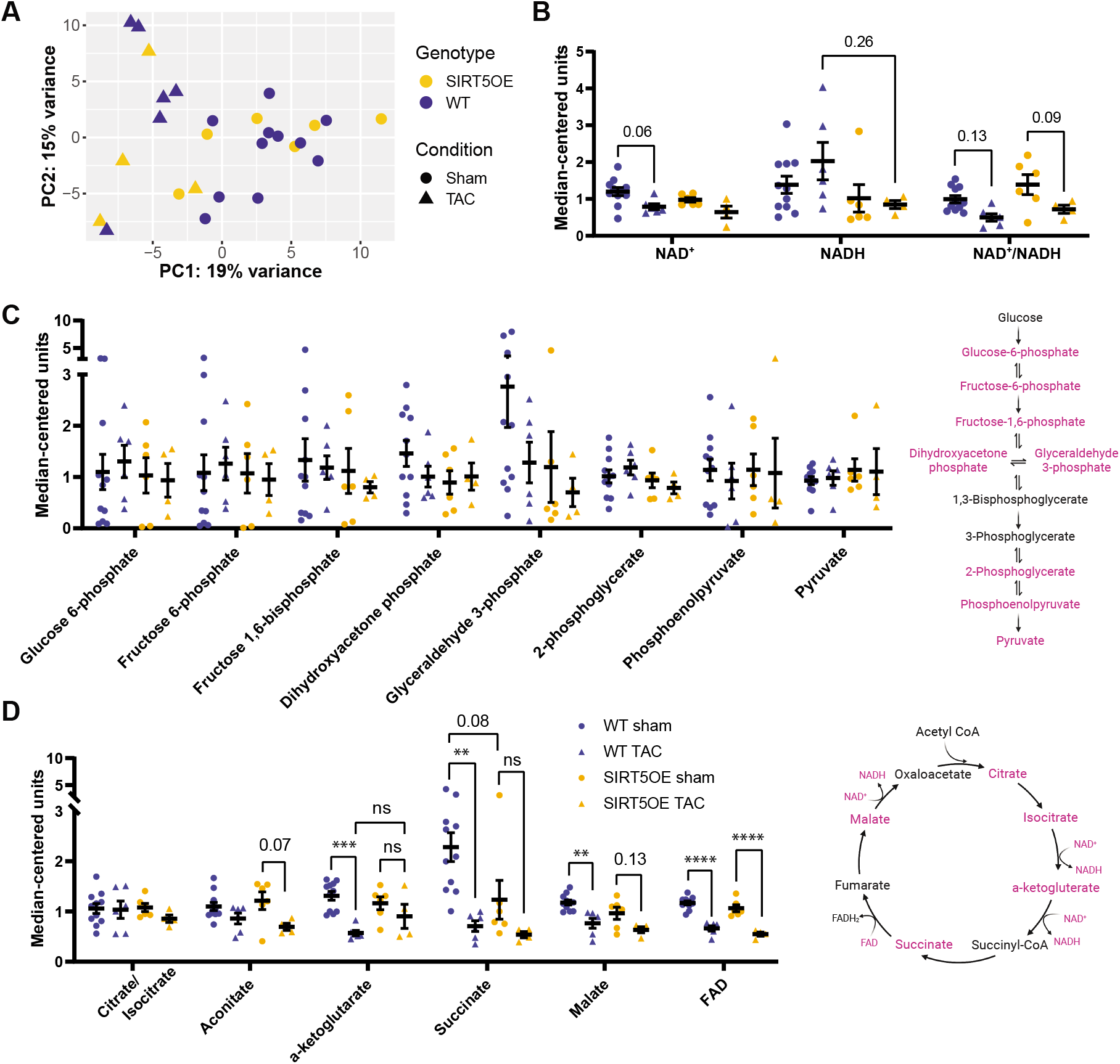
TAC condition is a strong determinant of the metabolomic landscape. A, PCA of the metabolomics data across all groups (WT sham (n=11), WT TAC (n=6), SIRT5OE sham (n=6) and SIRT5OE TAC (n=4)). B-D, metabolites of NAD+ (B), glycolysis (C), and the TCA cycle (D) plotted using median-centered values. Metabolites highlighted in magenta in the schematics represent measured metabolites.

Like other sirtuins, SIRT5 enzymatic activity is dependent on NAD^+^, an essential cofactor in oxidative phosphorylation and redox biology. Two-way ANOVA identified a significant TAC effect on NAD^+^ levels, and multiple comparisons testing showed a trend towards decreased NAD^+^ after TAC in WT mice (Figure 6B). WT hearts exhibited a larger range of NADH measurements, and genotype was a significant influence on NADH levels. In line with previous studies, we also found that the overall NAD^+^/NADH ratio was significantly decreased by TAC, and individual comparisons between sham and TAC groups revealed non-significant decreases (Supplemental Table 1) (32). These results suggest that an increase in oxidative stress occurs in the heart during TAC. Consistently, RNA-seq gene ontology analysis showed that both HIF1α signaling and NRF2-mediated oxidative stress response were significantly enriched in both sham vs TAC comparisons (Table 2).

Increased glucose usage and glycolytic flux are commonly observed during HF (33). We assessed metabolic intermediates in glycolysis and the citric acid (TCA) cycle to identify trends in these pathways. Glycolytic metabolites did not show any significant alterations (Figure 6C). In contrast, aside from citrate/isocitrate, levels of most TCA cycle metabolites were significantly altered by TAC (Supplemental table 1). α-ketoglutarate, succinate, and malate all significantly decreased in WT hearts after TAC, whereas SIRT5OE TAC animals showed non-significant trends towards decreases in these metabolites compared to sham (Figure 6D). Taken together, transcriptomics and metabolomics reflect a primarily TAC-driven, rather than genotype-associated, shift in the metabolic landscape in both genotypes following TAC.

### Cardiac fibrosis is suppressed in the SIRT5OE mice after TAC

Aside from metabolic pathways, another transcriptional signature identified by gene ontology analysis in TAC animals was extracellular matrix remodeling and fibrosis (Table 2, Figure 4D-F). GSEA showed that WT TAC samples were significantly enriched with genes mapping to TGFβ1 signaling, a cytokine central to the fibrotic response and the ECM, macromolecules secreted by fibroblasts essential to providing support to, and maintaining structure for, the myocardium (Figure 7A) (24). Hepatic Fibrosis Signaling was also highly significant in the IPA analysis and includes signaling events that occur during fibroblast activation, a process in which resident fibroblasts differentiate into myofibroblasts (Table 2). Fibroblast activation is essential to the adaptive hypertrophy process, as myofibroblasts gain the ability to contract surrounding tissue and remodel the ECM (24, 34). However, sustained myofibroblast activity results in excessive cardiac remodeling and is also a major contributor to LV dysfunction (34). Thus, according to GSEA and IPA gene ontology analysis, SIRT5 overexpression dampened the fibrotic signaling response following TAC.

**Figure 7.**
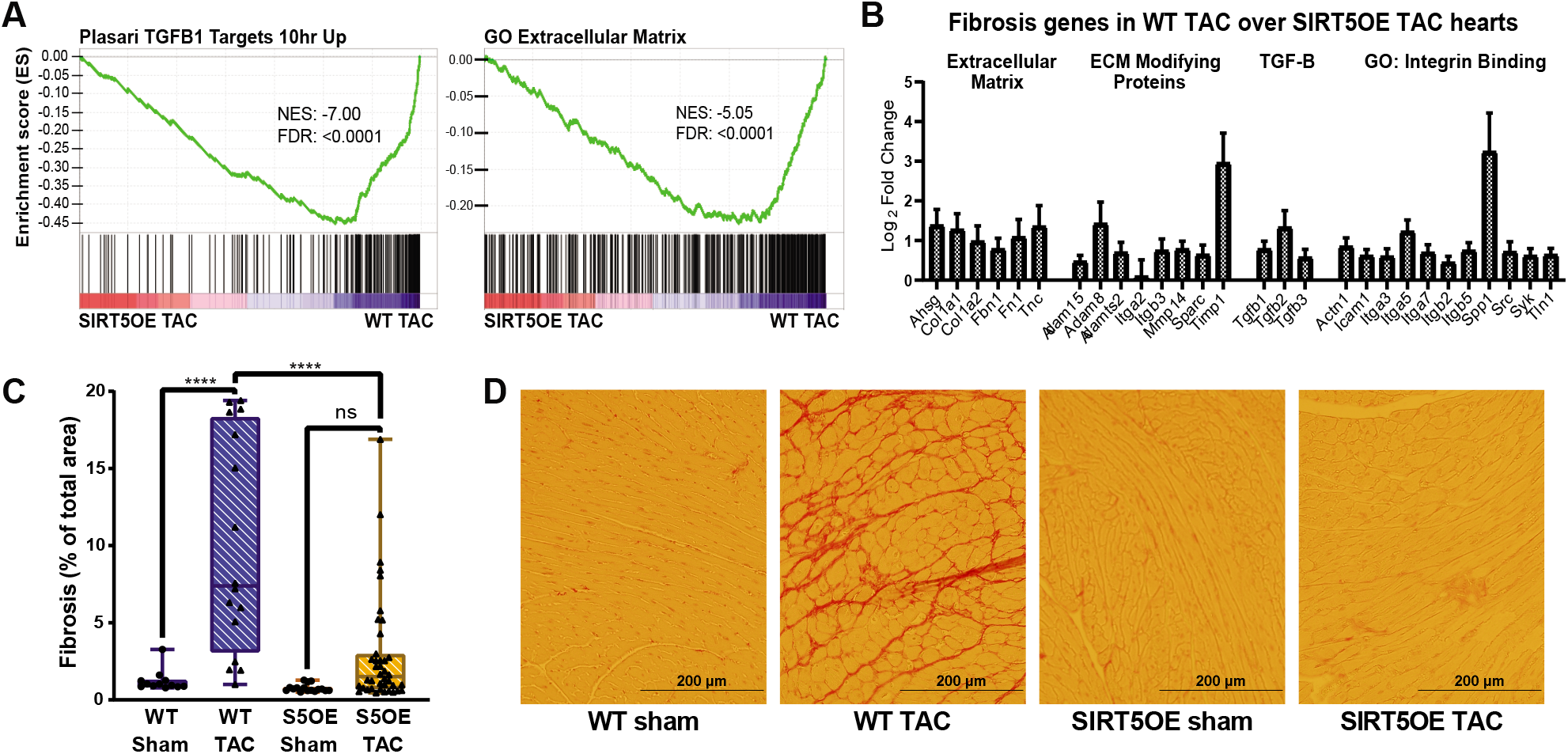
SIRT5OE protects against TAC-induced fibrosis. A, GSEA of SIRT5OE TAC compared to WT TAC RNA-seq samples. NES: normalized enrichment score. FDR: false discovery rate. B, a panel of select genes involved in fibrosis; all are significantly differentially expressed (FDR < 0.05) in SIRT5OE TAC compared to WT TAC hearts. Error bars represent standard error of the mean (SEM) of each gene. C, quantification of the amount of fibrosis in each heart sample, expressed in percentage ((WT sham (n=3), WT TAC (n=4), SIRT5OE sham (n=4) and SIRT5OE TAC (n=10)). Two-way ANOVA interaction term: p-value = 0.0012. D, representative images of heart sections stained with Picrosirius red for collagen.

In addition to promoting fibroblast activation, TGFβ signaling also induces expression of ECM components (*Col1a1, Col1a2, Fbn1, Tnc*), ECM modifying genes (*Timp1, Adam8, Adam15, Mmp14*), TGFβ cytokines, and cell adhesion genes (*Icam1, Itga3, Itgb2, Spp1*) (Figure 7B) (24, 35). mRNA levels of these genes were significantly higher in the WT hearts compared to SIRT5OE hearts after TAC. Thus, SIRT5 overexpression suppresses pressure overload induced TGFβ related fibrosis signaling. These differences in mRNA expression were also reflected in tissue histology. Staining for collagen using picrosirius red showed a dramatic increase in ECM accumulation in WT TAC but not SIRT5OE TAC tissue (Figure 7C, D). We conclude that SIRT5 overexpression blunts TAC-inducted cardiac fibrosis, concomitant with maintaining cardiac function.

## Discussion

These data demonstrate that SIRT5 is a limiting factor in maintaining cardiac structure and function during chronic pressure overload. Supra-physiological SIRT5 expression protects against HF-associated phenotypes, in particular systolic dysfunction and fibrosis. Elevated SIRT5 levels promoted maintenance of cardiac contractile function after 4 weeks of pressure overload, at which point WT mice had developed systolic dysfunction, characterized by decreased EF, ventricular dilation, remodeling and fibrosis. Consistently, RNA-seq analysis revealed that key cardiac disease-related signaling pathways, including TGFβ, IL6, Renin-Angiotensin, and NFAT, were blunted in SIRT5OE TAC mice. These data indicate a cardioprotective role of SIRT5 in response to stress, and are consistent with several previous studies showing that SIRT5KO mice are more susceptible to damage after cardiac stress (12, 16–18).

At least three possible non-mutually exclusive models may explain how SIRT5 confers protection against the deleterious consequences of TAC-induced pressure overload. First, SIRT5-mediated regulation of specific metabolic pathways is an attractive model, as metabolic dysfunction is both a driver and a consequence of hypertrophy and HF (30). A variety of proteomics and SIRT5 activity studies have shown that SIRT5-dependent regulation of its targets can be highly context-dependent (36). Studies conducted using SIRT5KO mice and cell culture models have indicated that SIRT5 promotes fatty acid oxidation by desuccinylating ECHA (12). As noted previously, the heart shifts from fatty acid oxidation to increased glucose consumption upon hypertrophy and failure. In our studies, WT TAC mice exhibited transcriptional downregulation of genes involved in fatty acid catabolism and oxidative phosphorylation, with concurrent upregulation of glycolytic genes, compared to SIRT5OE TAC mice. However, these transcriptional signatures were not reflected in changes in ETC protein expression or electron flow through the ETC. Notably, in an ischemia-reperfusion model, SIRT5KOs exhibited a larger infarct size, and inhibition of SDH reduced this infarct size in both WT and SIRT5KO mice (16). Furthermore, succinate injection is sufficient to induce cardiomyocyte hypertrophy, while inhibition of SDH results in improved recovery following ischemia-reperfusion (37, 38). Our results and those of others have revealed that SIRT5 modulates SDH activity, in a cell- and context-dependent fashion, implying that SIRT5OE hearts may have altered SDH activity, which could contribute to their cardioprotection (8, 36, 39). In this regard, metabolomics revealed a nearly significant (p = 0.08) increase in succinate levels in the WT sham hearts compared to SIRT5OE sham hearts, differences that were blunted following TAC (Figure 6D). Therefore, overexpression of SIRT5 might protect against hypertrophy and HF through regulating levels of succinate, and/or other key metabolites.

ROS management may represent another target of SIRT5 in the context of SIRT5-mediated cardioprotection. Mitochondria represent the major source of cellular ROS production. Increased ROS production due to pathological stimuli, including cytokine signaling or mechanical stretching, can stimulate cardiomyocyte hypertrophy, ventricular dilation, and contractile dysfunction (40). Rat myoblasts treated with H2O2 show increased apoptosis and decreased SIRT5 levels; in contrast, we did not observe changes in SIRT5 transcript or protein levels after TAC (Supplemental Figure 3A, B). In rat myoblasts, SIRT5 overexpression protected against H2O2-driven apoptosis through interactions with BCL-XL, an anti-apoptotic member of the Bcl-2 family (41). Likewise, in other cell types, SIRT5 has been shown to reduce cellular ROS levels via desuccinylation-mediated activation of SOD1 and promote NADPH production by desuccinylation and deglutarylation of IDH2 and G6PD, respectively (36). Elevated cellular ROS induces expression of a specific set of genes to detoxify ROS, an effect mediated in part through the NRF2 transcription factor (42). NRF2-mediated oxidative stress response was significantly enriched in both WT and SIRT5OE RNA-seq comparisons but blunted in the latter group (Table 2). Future studies are needed to determine whether this effect is a driver or a consequence of attenuated cardiac dysfunction in SIRT5OE mice.

Second, immune-related pathways were among the most significantly altered by SIRT5OE in the context of TAC and may be relevant for cardioprotection conferred by SIRT5 overexpression (Fig 5D). Early immune infiltration after TAC is necessary for the transition from ventricular hypertrophy to dilation during pressure overload (43, 44). Blocking this inflammatory response alleviates late LV remodeling and dysfunction, cardiac fibrosis, and T cell expansion (45–47). Similarly, in our studies, both genotypes developed significant hypertrophy after 4 weeks of TAC, but SIRT5OE mice were protected against subsequent ventricular dilation and dysfunction (Fig 2C-D; 3A-C). Although current published literature on SIRT5 in the immune system is limited, SIRT5 might protect against TAC by dampening the pro-inflammatory response in macrophages, where SIRT5 has been shown to desuccinylate and activate PKM2, thereby blocking LPS-driven macrophage activation (48). Likewise, malonylation of GAPDH promotes TNFα translation and macrophage activation, although SIRT5’s role in this process is uncharacterized (49). Consistently, single-cell RNA-seq data from TAC mice reveals that SIRT5 is expressed in macrophage and T-cell populations, in addition to cardiomyocytes and other cell types in the heart (Supplemental Figure 3B,C) (47). In this regard, two studies from the Hirschey group showed that while constitutive whole-body SIRT5KO mice displayed increased mortality after TAC, cardiomyocyte-specific SIRT5KOs and controls responded to TAC similarly, raising the possibility of cardiomyocyte-non-autonomous functions of SIRT5 in response to TAC (17, 18).

Third, SIRT5 might regulate cardiac fibrosis, a process initiated and sustained through a variety of signaling and mechanical factors, including inflammatory cytokines (*e.g.,* TGFβ, Angiotensin II, TNF*α*), neuroendocrine factors, and ventricular wall stretch (24). These factors signal cardiac fibroblasts to activate, migrate, proliferate, and differentiate into myofibroblasts. In addition to ECM secretion and remodeling, activated fibroblasts exert protective effects through suppressing proinflammatory phenotypes and even promote the hypertrophic response through secretion of critical cytokines (50, 51). However, chronic myofibroblast activity inhibits oxygen and nutrient delivery, resulting in myocyte atrophy and death, ultimately leading to ventricular dilation and dysfunction (24, 34). The fibrotic response following TAC represents one of the most pronounced differences between WT and SIRT5OE mice. SIRT5 may play a role in regulating cytokine-mediated activation of fibrosis, either in immune cells where cytokines originate, in triggering fibroblast activation, and/or in myofibroblasts themselves. Further studies are needed to determine (1) how long SIRT5 overexpression protects mice from dilation and contractile failure after TAC and (2) in which cell and tissue type(s) SIRT5 functions to confer cardioprotection.

In summary, we have shown that increased expression of the deacylase SIRT5 is sufficient to confer improved maintenance of cardiac function and decreased fibrosis during pressure overload. Sirtuin activity is amenable to enhancement by allosteric activators and NAD+ precursors, and we found that NADH levels and NAD^+^/NADH ratios were significantly impacted by SIRT5 overexpression (Figure 6B, Supplemental table 1) (52, 53). Our results hint that SIRT5 might represent an attractive novel target in cardiac fibrosis, a condition that afflicts millions in the US and worldwide, but which currently lacks mechanism-based therapies.

## Methods

### Generation of SIRT5OE mice

Mouse ES cell clone 5-11C was derived from V6.5 ES cells (Pittman et al., 1998). ES cells were cultured in high-glucose Dulbecco’s minimal essential medium supplemented with 15% fetal bovine serum, 1 μM 2-mercaptoethanol, 4 mM glutamine, 50 IU of penicillin per ml, 50 ug of streptomycin per ml, and 1,000 U of recombinant leukemia inhibitory factor (ESGRO; Millipore) on mitotically inactivated feeder cells as described (Hughes and Saunders, 2011). Chromosome counting indicated that 90% of twenty counted chromosome spreads contained an euploid chromosome number. ES cells were microinjected into 96 blastocysts (Bradley, 1987) obtained from the mating of albino mice (C57BL/6BrdCrHsd-*Tyr^c^*, Envigo). Of twenty-one pups born, twenty were ES cell-mouse chimeras and eighteen of these were male chimeras with at least 70% contribution from 5-11C ES cells, as judged by coat color contribution. Sox2-Cre (JAX 008454) were mated with mice containing the SIRT5OE cassette to delete the loxP-STOP-loxP site to generate a global SIRT5OE transgenic mouse.

### Immunoblotting

Mice were anesthetized with isoflurane and immediately sacrificed by cervical dislocation. Heart tissue samples were flash frozen, pulverized, and resuspended in Laemmli sample butter (62.5mM Tris pH6.8, 2% SDS, 10% glycerol, 5% β-mercaptoethanol, 1% bromophenol blue). Samples were sonicated for 30 seconds on ice, then spun down at 4°C for 30 minutes at 15,000xg, and the supernatant was collected for protein quantification using a DC Protein Assay (Bio-Rad #5000112). Samples were boiled, resolved by SDS-PAGE, and transferred to PVDF membrane overnight at 4°C using a Bio-Rad Criterion system. Membranes were stained with Ponceau S to assess protein loading, and then blocked using 5% milk in TBST (TBS containing 0.1% Tween-20) at room temperature. Primary antibodies were diluted in 5% BSA in TBST and incubated with the blot overnight at 4°C on a rocker. Secondary incubation was performed at room temperature for 1 hour using either mouse or rabbit secondary antibodies (Jackson ImmunoResearch 115-035-062 or 111-035-045) diluted 1:10000 in 5% milk in TBST. For detection, blots were immersed in a western chemiluminescent HRP substrate (Millipore P90720) and imaged using the GE ImageQuant LAS 4000. Primary antibodies used in this study: SIRT5 (Cell Signaling #8782), Succinyl-lysine (PTMBiolabs #PTM-419), Malonyl-lysine (PTMBiolabs #PTM-901), SOD2 (Santa Cruz #30080), GAPDH (Cell Signaling #5174P), Rodent OXPHOS (Abcam #ab110413), Sarcomeric Actin (Sigma # A2172-100UL), Tubulin (Santa Cruz # SC-23948).

### TAC procedure

TAC was performed at the University of Michigan Physiology and Phenotyping Core. Mice were anesthetized using Isoflurane. Carprofen use administered pre-emptively and for 48 hours, then as needed. The animals were intubated and ventilated. The aortic arch was isolated by entering the extrapleural space above the first rib, and the transverse aorta was isolated between the right and left carotid arteries. A 7-0 or similar size nylon suture ligature was tied around the transverse aorta against a 27-gauge needle to produce a 60-70% constriction after the removal of the needle. The incision was closed, and the left-sided pneumothorax (if present) was evacuated.

### Human heart tissue procurement

Ventricular myocardial tissue from patients with advanced heart failure at the University of Michigan was collected at the time of cardiac transplantation following obtaining of written informed consent. Nonfailing ventricular myocardial tissue was collected from unmatched donor hearts from the University of Michigan. Co-morbidities in the donors that may have precluded use of their hearts for cardiac transplantation included age, hypertension, diabetes, minor coronary artery disease, alcohol or tobacco use. Prior to tissue retrieval, all hearts were perfused with ice-cold cardioplegia. Samples from each heart were snap frozen in liquid N2 at the time of arrival and stored at −80°C. This study was approved by the institutional IRB.

### Echocardiography

Induction of anesthesia was performed in an enclosed container filled with 5% isoflurane. After induction, the mice were placed on a warming pad to maintain body temperature. 1%-1.5% isoflurane was supplied via a nose cone. Hair was removed from the upper abdominal and thoracic area with depilatory cream. ECG was monitored via non-invasive resting ECG electrodes. Transthoracic echocardiography was performed in the supine or left lateral position. Two-dimensional, M-mode, Doppler and tissue Doppler echocardiographic images were recorded using a Visual Sonics’ Vevo 2100 high resolution *in vivo* micro-imaging system. LV ejection fraction was measured from the two-dimensional long axis view. Systolic and diastolic dimensions and wall thickness were measured by M-mode in the parasternal short axis view at the level of the papillary muscles. Fractional shortening and ejection fraction were also calculated from the M-mode parasternal short axis view. Diastolic function was assessed by conventional pulsed-wave spectral Doppler analysis of mitral valve inflow patterns (early [E] and late [A] filling waves). Doppler tissue imaging (DTI) was used to measure the early (Ea) diastolic tissue velocities of the septal and lateral annuluses of the mitral valve in the apical 4-chamber view.

### Mitochondrial isolation

Mitochondrial isolation was performed as previously described (37). In brief, mice were anesthetized with isoflurane and immediately sacrificed by cervical dislocation. After cardiac harvest, ventricles were minced in ice-cold MIB (210 mM D-Mannitol, 70 mM sucrose, 5 mM MgCl_2_, 10 mM KH_2_PO_4_, 5 mM HEPES, 1 mM EGTA, 0.2% fatty acid-free BSA), mechanically homogenized using a Fisher Scientific PowerGen 125, and centrifuged at 800x*g* then 8000x*g* for 10 minutes at 4°C. The final pellets, containing mitochondria, were resuspended in MAS (220 mM D-Mannitol, 70 mM sucrose, 5 mM MgCl_2_, 10 mM KH_2_PO_4_, 2 mM HEPES, 1 mM EGTA, 0.1% fatty acid-free BSA). Mitochondria were quantified using DC quantification assy. Both MIB and MAS pH were adjusted to 7.2 with KOH and stored at 4°C before use.

### Agilent/Seahorse assay

To assess the functionality of isolated mitochondria from WT and SIRT5OE hearts, the electron flow assay was performed using the XFe96 Extracellular Flux Analyzer (Agilent Technologies, Santa Clara, CA) per the manufacturer’s instructions. Briefly, 1μg of isolated mitochondria (from either WT or SIRT5OE mice) in 25μl of mitochondrial assay solution (MAS), supplemented with 10mM pyruvate, 2mM malate, and 4μM FCCP, were plated per well of a 96-well Seahorse microplate on ice. The microplate was then transferred to a centrifuge equipped with a swinging bucket microplate adaptor, and spun at 2000g for 20 minutes at 4°C. After centrifugation, 155μL of pre-warmed (37°C) MAS, supplemented with 10mM pyruvate, 2mM malate, and 4μM FCCP, was added to each well. Mitochondrial oxygen consumption rates (OCR) were then measured, under initial conditions and following addition of 2μM rotenone (Sigma #R8875), 10mM succinate (Sigma #S9515), 4μM antimycin A (Sigma #A8674), and 10mM ascorbate (Sigma #A7506) plus 100μM TMPD (Sigma #T3134).

### Metabolomics sample preparation

Mice were anesthetized with isoflurane and immediately sacrificed by cervical dislocation. Heart tissue samples were immediately flash frozen, pulverized to powder, vortexed, and centrifuged for 10 min at 13000 x g, 4°C. Then, 1 ml of the supernatant was aspirated from each tube, transferred to a tightly capped sample tube, and stored at −80°C until analysis.

### Metabolomics MS/MS system

Samples were run on an Agilent 1290 Infinity II LC −6470 Triple Quadrupole (QqQ) tandem mass spectrometer (MS/MS) system with the following parameters: Agilent Technologies Triple Quad 6470 LC-MS/MS system consists of the 1290 Infinity II LC Flexible Pump (Quaternary Pump), the 1290 Infinity II Multisampler, the 1290 Infinity II Multicolumn Thermostat with 6 port valve and the 6470 triple quad mass spectrometer. Agilent Masshunter Workstation Software LC/MS Data Acquisition for 6400 Series Triple Quadrupole MS with Version B.08.02 was used for compound optimization, calibration, and data acquisition.

### Metabolomics liquid chromatography

2μL of sample was injected into an Agilent ZORBAX RRHD Extend-C18 column (2.1 × 150 mm, 1.8 um) with ZORBAX Extend Fast Guards. The LC gradient profile is as follows, solvent conditions below. 0.25 ml/min, 0-2.5 min, 100% A; 2.5-7.5 min, 80% A and 20% B; 7.5min-13 min 55% A and 45% B; 13min-24 min, 1% A and 99% B; 24min-27min, 1% A and 99% C; 27min-27.5min, 1% A and 99% C; at 0.8 ml/min, 27.5-31.5 min, 1% A and 99% C; at 0.6 ml/min, 31.5-32.25min, 1% A and 99% C; at 0.4 ml/min, 32.25-39.9 min, 100% A; at 0.25 ml/min, 40 min, 100% A. Column temp is kept at 35 °C, samples are at 4 °C.

### LC Solvents

Solvent A is 97% water and 3% methanol 15 mM acetic acid and 10 mM tributylamine at pH of 5. Solvent B is 15 mM acetic acid and 10 mM tributylamine in methanol. Washing Solvent C is acetonitrile. LC system seal washing solvent 90% water and 10% isopropanol, needle wash solvent 75% methanol, 25% water. Solvents were purchased from the following vendors: GC-grade Tributylamine 99% (ACROS ORGANICS), LC/MS grade acetic acid Optima (Fisher Chemical), InfinityLab Deactivator additive, ESI–L Low Concentration Tuning mix (Agilent Technologies), LC-MS grade solvents of water, and acetonitrile, methanol (Millipore), isopropanol (Fisher Chemical).

### Metabolomics mass spectrometry

6470 Triple Quad MS is calibrated with the Agilent ESI-L Low Concentration Tuning mix. Source parameters: Gas temp 150 °C, Gas flow 10 l/min, Nebulizer 45 psi, Sheath gas temp 325 °C, Sheath gas flow 12 l/min, Capillary −2000 V, Delta EMV −200 V. Dynamic MRM scan type is used with 0.07 min peak width, acquisition time is 24 min. Delta retention time of plus and minus 1 min, fragmentor of 40 eV and cell accelerator of 5 eV are incorporated in the method. The MassHunter Metabolomics Dynamic MRM Database and Method was used for target identification. Key parameters of AJS ESI were: Gas Temp: 150 °C, Gas Flow 13 l/min, Nebulizer 45 psi, Sheath Gas Temp 325 °C, Sheath Gas Flow 12 l/min, Capillary 2000 V, Nozzle 500 V. Detector Delta EMV(-) 200.

### Metabolomics data analysis

The QqQ data were pre-processed with Agilent MassHunter Workstation QqQ Quantitative Analysis Software (B0700). Each metabolite abundance level in each sample was divided by the median of all abundance levels across all samples for proper comparisons, statistical analyses, and visualizations among metabolites. A total of 230 metabolites were measured. Statistical significance was calculated by a two-tailed t-test with a significance threshold level of 0.05. Median-centered values for all metabolites measured are recorded in Supplemental Table 2.

### Tissue preparation for RNA extraction

Heart tissue samples were flash frozen, pulverized, resuspended in TRIzol and homogenized using a power homogenizer (Fisher Scientific PowerGen 125). RNA was extracted using RNeasy Mini Kit (Qiagen 741-4) per the manufacturer’s instructions.

### qRT-PCR

cDNA was generated from RNA using SuperScript III Reverse Transcriptase (Invitrogen 18080044) per the manufacturer’s instructions. TaqMan Gene Expression Assays (Applied Biosystems) for single-tube assays were used for all qRT-PCR experiments. Reactions were carried out as per the manufacturer’s methods, using TaqMan Gene Expression Assays as listed with the associated TaqMan Assay ID: *Myh6* (Mm00440359_m1), *Acta1* (Mm00808218_g1), *Nppa* (Mm01255747_g1), *Myh7* (Mm00600555_m1).

### RNA-sequencing bioinformatic analysis

RNA libraries were prepared using a standard Illumina protocol. Base-calling was performed on an Illumina HiSeq4000. Raw fastq files were analyzed using FastQC for quality control. Transcripts were aligned to mm ensemble cDNA release 101 using kallisto (v0.46.0) and counted using Tximport (v1.18.0) (54–56). DESeq2 was used for calculating differential expression and results are documented in Supplemental Table 3 (57). Genes with a false discovery rate (FDR) was less than 0.05 was designated as differentially expressed between groups. Pathway analysis was performed using Ingenuity Pathway Analysis (IPA) and gene set enrichment analysis (GSEA) (28, 29). These data have been submitted to GEO, accession #XXXX.

### Fibrosis staining

Heart tissue was paraffin embedded, sectioned, and stained overnight with picrosirius red at the University of Michigan Dentistry Histology Core. Paraffin embedded heart section were de-paraffinized in xylene, rehydrated in 100% EtOH, 95% EtOH, and 70% EtOH and dH20. Sections were then fixed in 10% buffered formalin and stained in picric acid solution with 0.1% Sirius red, dehydrated in 80% EtOH, 95% EtOH, then 100% EtOH, cleared in xylene, and finally mounted with Permount. For each heart, eight images of left ventricle sections, 500 *μ*m x 400 *μ*m, were acquired under polarized bright-filed light for fibrosis detection. Collagen area fraction (CAF) of the total areas, 500 *μ*m x 400 *μ*m, were measured by ImageJ using the equation ((collagen area)/(total area)=CAF). 4 images with the highest CAF were used for statistical analysis.

### Statistics

Statistical analysis were conducted using Graphpad Prism, version 8. Multiple-group comparisons were analyzed using two-way ANOVA followed by Sidak’s correction for multiple comparisons. For analysis of two groups, Welch’s t-test was used. A p-value of less than 0.05 was considered statistically significant. Scatter plots include median and interquartile range markers. Significance markers: (*) p < 0.05, (**) p < 0.01, (***) p < 0.001, (****) p < 0.0001.

### Study approval

All mice were housed at the Biomedical Science Research Building (UM). All vertebrate animal experiments were approved by and performed in accordance with the regulations of the University Committee on Use and Care of Animals.

## Author contributions

AHG, RB, MES, SK, AA, LZ, NJD, MWM, SMD, CAL, ABS, and DBL performed experiments and/or analyzed data. SM and DAS provided the SIRT5OE mouse ES cells. AHG, ABS, and DBL interpreted data. AHG and DBL wrote the manuscript. AHG made the figures. DBL and ABS supervised overall design and study interpretation.

## Acknowledgements

The authors would like to thank the University of Michigan Transgenic Animal Model Core for assistance with generation of the SIRT5OE mouse strain; Steven Whitesall, Kimber Converso-Baran, and Dr. Daniel Michele (Physiology and Phenotyping Core) for TAC and echo services and helpful discussions; and the Dentistry Histology Core. Dr. Heiko Bugger is also acknowledged for helpful discussions. We would also like to thank Dr. Daniel Goldstein and the other members of the Guo dissertation committee, and members of the Lombard Lab for useful feedback. This project was supported by R01GM101171, R21ES032305 and the Glenn Foundation for Medical Research (GFMR) (DBL); T32AG000114 and T32GM113900 (AHG); R25GM086262 (RKB); R37AG028730 and GFMR (DAS); R01CA248160, R01CA244931 and UMCCC Core Grant: P30CA046592 (CAL). ABS was funded by the University of Michigan Frankel Cardiovascular Center, the University of Michigan McKay Grant, and the Taubman Emerging Scholars Program. Metabolomics studies performed at the University of Michigan were supported by DK097153, the Charles Woodson Research Fund, and the UM Pediatric Brain Tumor Initiative. NJD was supported by the Frankel Cardiovascular Center Summer Undergraduate Fellowship Program.

## Conflict of interest

DAS is a consultant to MetroBiotech, a company developing NAD^+^ boosters to treat rare diseases; a complete list of DAS activities is at https://sinclair.hms.harvard.edu/david-sinclairs-affiliations. CAL has received consulting fees from Astellas Pharmaceuticals and is an inventor on patents pertaining to Kras regulated metabolic pathways, redox control pathways in pancreatic cancer, and targeting the GOT1-pathway as a therapeutic approach. DBL reports ownership of the equivalent in voting stock or share of ABBV and GILD.

