## Supplemental Figures for "Sirtuin 5 levels are limiting in preserving cardiac function and suppressing fibrosis in response to pressure overload"

### Supplemental Figure 1

**A**

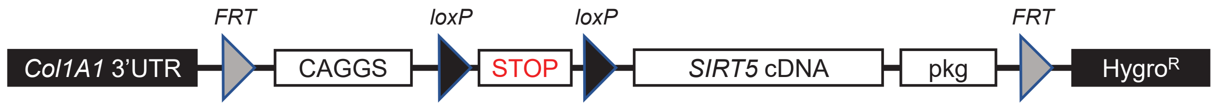


**B**

| WT male, SIRT5OE female | | | | | | |
|---|---|---|---|---|---|---|
| | **WT** | **SIRT5OE** | **Total** | | $\chi^2$ | **p-value** |
| **Males** | 73 | 77 | 150 | | 2.09 | 0.55 |
| **Females** | 74 | 61 | 135 | | | |
| **Total** | 147 | 138 | 285 | | | |

| SIRT5OE male, WT female | | | | | | |
|---|---|---|---|---|---|---|
| | **WT** | **OE** | **Total** | | $\chi^2$ | **p-value** |
| **Males** | 38 | 25 | 63 | | 5.07 | 0.17 |
| **Females** | 42 | 41 | 83 | | | |
| **Total** | 80 | 66 | 146 | | | |

| Combined | | | | | | |
|---|---|---|---|---|---|---|
| | **WT** | **SIRT5OE** | **Total** | | $\chi^2$ | **p-value** |
| **Males** | 111 | 102 | 213 | | 1.32 | 0.72 |
| **Females** | 116 | 102 | 218 | | | |
| **Total** | 227 | 204 | 431 | | | |

**Supplemental Figure 1. Generation of SIRT5OE mice.** A, cassette containing a constitutive CAGGS promoter, a transcriptional flox-STOP-flox, followed by the SIRT5 cDNA was inserted into the collagen1 A1 (Col1A1) 3'UTR by FLP recombination. B, number of male and female mice born per WT or SIRTOE litter.

### Supplemental Figure 2

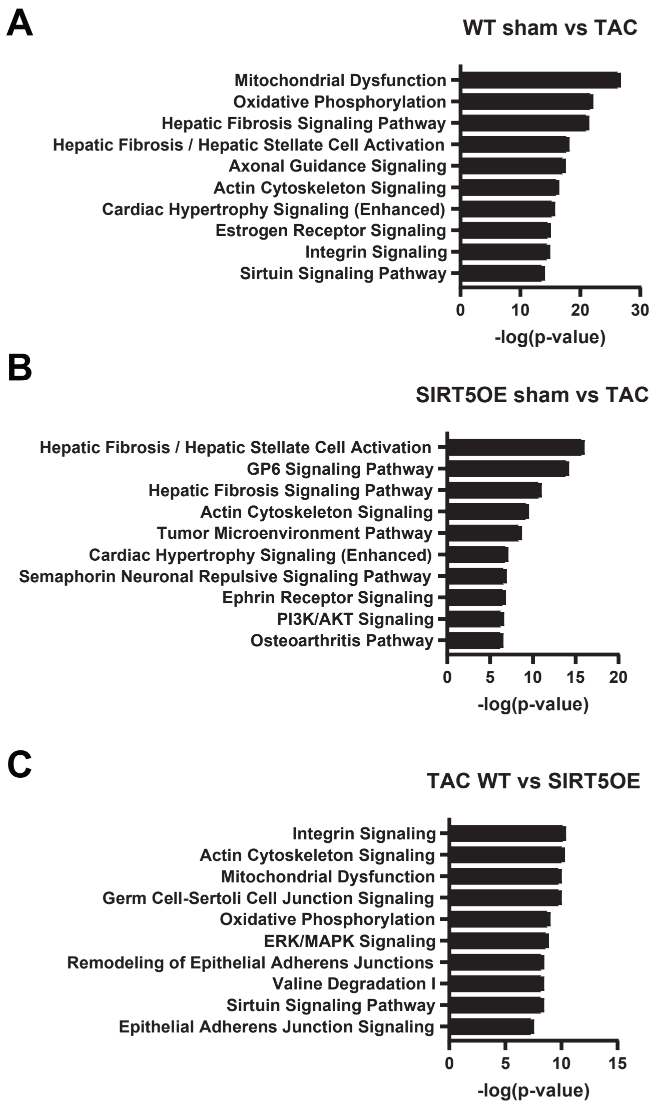


**Supplemental Figure 2.** Top 10 IPA pathways, sorted on significance, enriched by differentially expressed genes in each labelled comparison.

Supplemental Figure 3

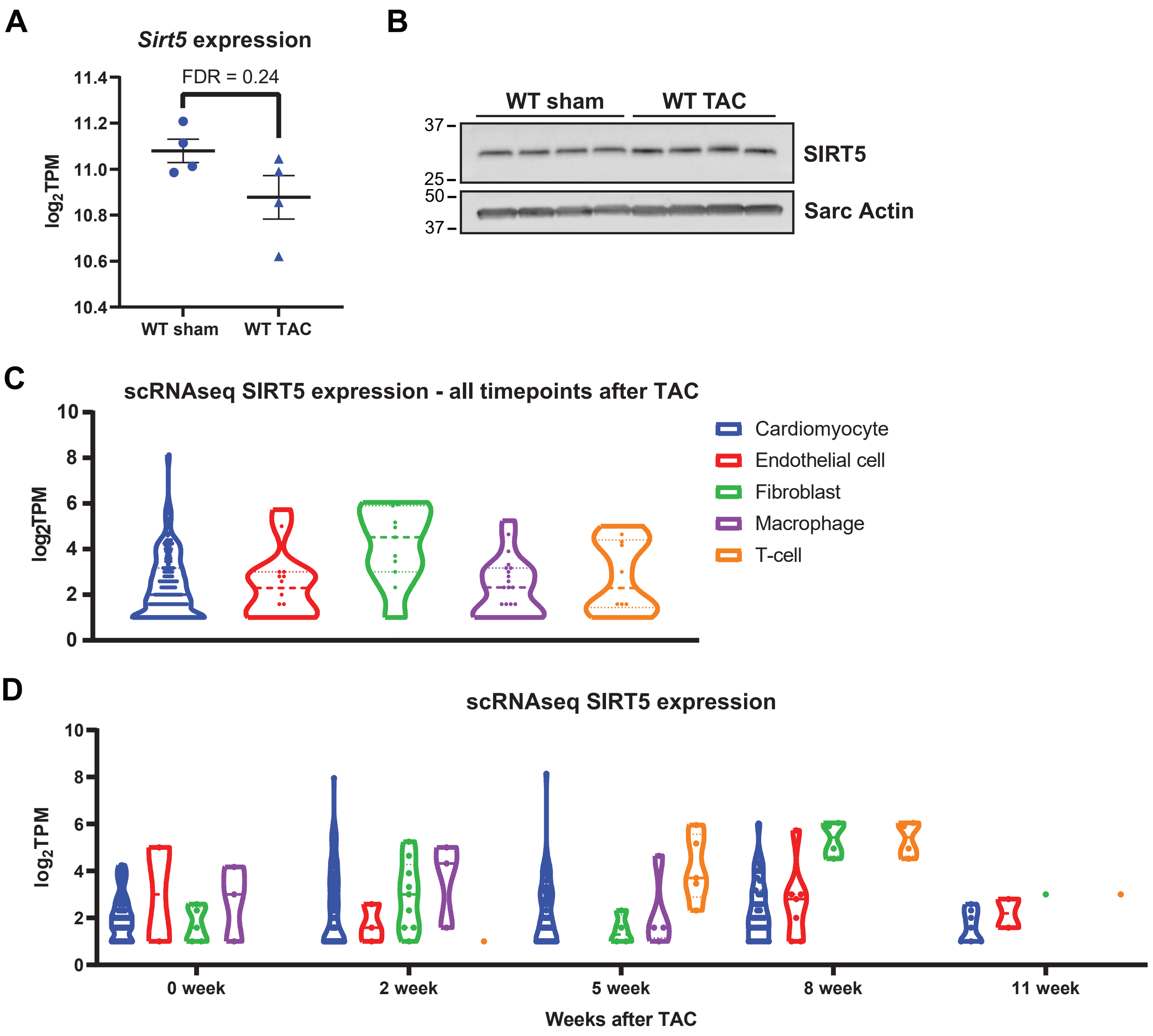


**Supplemental Figure 3. SIRT5 expression in bulk heart tissue and specific cardiac cell populations.** A, Sirt5 expression in bulk heart WT sham and WT TAC samples. B, SIRT5 protein levels in WT sham and TAC heart lysates. C, Sirt5 expression in five major cardiac cell types using single-cell data generated by Ren et. al. Only cells with at least 2 reads of Sirt5 were graphed. D, Sirt5 expression in single-cell RNA-seq across 5 different time-points through 11 weeks of pressure overload.

| **Supplemental Table 1. Two-way ANOVA analyses of metabolites in Figure 6B-D** | | | |
|---|---|---|---|
| | **Genotype Effect** | **TAC Effect** | **Interaction Effect** |
| **$NAD^+$** | 0.136 | 0.005* | 0.740 |
| **NADH** | 0.044* | 0.522 | 0.275 |
| **$NAD^+$/NADH** | $0.074^+$ | 0.002* | 0.599 |
| **Glucose 6-phosphate** | 0.584 | 0.888 | 0.705 |
| **Fructose 6-phosphate** | 0.694 | 0.945 | 0.712 |
| **Fructose 1,6-bisphosphate** | 0.504 | 0.598 | 0.850 |
| **Dihydroxyacetone phosphate** | 0.326 | 0.555 | 0.321 |
| **Glyceraldehyde 3-phosphate** | 0.197 | 0.235 | 0.548 |
| **2-phosphoglycerate** | 0.133 | 0.940 | 0.306 |
| **Phosphoenolpyruvate** | 0.825 | 0.696 | 0.832 |
| **Pyruvate** | 0.399 | 0.962 | 0.826 |
| **Citrate/Isocitrate** | 0.532 | 0.345 | 0.441 |
| **Aconitate** | 0.840 | 0.005* | 0.263 |
| **a-ketoglutarate** | 0.460 | <0.001* | $0.065^+$ |
| **Succinate** | $0.069^+$ | 0.002* | 0.183 |
| **Malate** | $0.058^+$ | <0.001* | 0.638 |
| **FAD** | 0.049* | <0.001* | 0.940 |

**Supplemental Table Legends**

**Supplemental Table 1.** Two-way ANOVA analyses results of all metabolites plotted in Figure 6B-D. Significance markers: (*) $p < 0.05$, (+) $p < 0.1$.

**Supplemental Table 2.** Median-centered data from LC-MS/MS metabolite profiling in WT sham, WT TAC, SIRT5OE sham, and SIRT5OE TAC hearts. P-value was calculated using a two-tailed t-test.

**Supplemental Table 3.** DESeq2 output of differential gene expression of the RNA-seq data. Samples in each comparison are labelled on the spreadsheet name. Padj (p-value adjusted for multiple comparisons) were used to determine significance.
